# Abrogation of Stem Loop Binding Protein (Slbp) function leads to a failure of cells to transition from proliferation to differentiation, retinal coloboma and midline axon guidance deficits

**DOI:** 10.1101/464123

**Authors:** Kate Turner, Jacqueline Hoyle, Leonardo E Valdivia, Kara Cerveny, Wendy Hart, Maryam Mangoli, Robert Geisler, Michele Rees, Corinne Houart, Richard J. Poole, Stephen W Wilson, Gaia Gestri

**Author notes:** These authors contributed equally to the work (Kate Turner, Jacqueline Hoyle).

## Abstract

Through forward genetic screening for mutations affecting visual system development, we identified prominent coloboma and cell-autonomous retinal neuron differentiation, lamination and retinal axon projection defects in *eisspalte* (*ele*) mutant zebrafish. Additional axonal deficits were present, most notably at midline axon commissures. Genetic mapping and cloning of the *ele* mutation showed that the affected gene is *slbp*, which encodes a conserved RNA stem-loop binding protein involved in replication dependent histone mRNA metabolism. Cells throughout the central nervous system remained in the cell cycle in *ele* mutant embryos at stages when, and locations where, post-mitotic cells have differentiated in wild-type siblings. Indeed, RNAseq analysis showed down-regulation of many genes associated with neuronal differentiation. This was coincident with changes in the levels and spatial localisation of expression of various genes implicated, for instance, in axon guidance, that likely underlie specific *ele* phenotypes. These results suggest that many of the cell and tissue specific phenotypes in *ele* mutant embryos are secondary to altered expression of modules of developmental regulatory genes that characterise, or promote transitions in, cell state and require the correct function of Slbp-dependent histone and chromatin regulatory genes.

**Author Summary:** Congenital deficits of eye formation are common in humans and to help understand the genetic basic of such conditions, we are studying zebrafish with comparable eye defects. We identified defects in both the shaping of the eye and in its connections to the brain in *eisspalte* mutant fish. Further analyses revealed additional deficits in the brain, most notably a severe reduction in neurons and their connections. We find that this is due to an inability of the cells that generate neurons to transition from proliferation to neuronal differentiation. By using a sequencing approach to compare mutant embryos to their normal siblings, we identified the affected gene as *slbp*, which encodes a protein that binds the mRNAs of other genes important for cell proliferation. This sequencing approach revealed the full extent of changes in gene expression in the mutant, helping us to better understand why the nervous system defects occur. Our study suggests that in the absence of Slbp function, cells lose the ability to transition from the proliferative to the differentiated state and this leads to additional defects in the eyes and brain.

## Introduction

Mutations in a wide variety of genes are known to lead to congenital abnormalities of eye formation [1,2]. Some of these genes, such as *pax6* and *rx3*, show temporally and spatially restricted expression within developing visual system structures and consequently, *a priori*, are obvious candidates for roles in eye formation [3]. However, other genes, such as *hdac1* [4] and *yap* [5], are more ubiquitously expressed and consequently visual system specific phenotypes observed upon aberrant gene function are not so easily explained. Forward genetic screens in animal models provide a relatively unbiased approach to identify the full spectrum of genes involved in specific developmental processes, as the initial selection is based upon phenotypes of interest [6]. To this end, we have been using a forward genetic approach in which we screen existing and new zebrafish lines carrying randomly induced mutations for phenotypes affecting visual system development.

In this study, we observed that in *eisspalte* (*ele*) mutants, the ventro-nasal and ventro-temporal lips of the forming eye cup fail to fuse, leading to prominent retinal coloboma. The *eisspalte* phenotype was originally identified on the basis of aberrant morphogenesis of the midbrain/hindbrain boundary [7] but the affected gene had not been identified. Using both traditional mapping approaches and a novel mapping-by-sequencing approach based on the variant discovery mapping Cloudmap pipeline [8,9], we find that the *eisspalte* mutation lies within the *slbp* gene. This is consistent with a description of retinal defects in another *slbp^rw440^* mutant allele [10].

Slbp is a stem loop RNA-binding protein required for all aspects of replication dependent histone mRNA metabolism. Replication-dependent histone genes, which are predominantly expressed during S-phase in proliferating cells, are intron-less and encode non-polyadenylated pre-mRNAs that are processed by an unusual mechanism that requires two cis-acting elements in their 3’ untranslated regions (UTR) referred to as the stem loop (SL) and the histone downstream element (HDE). Slbp binds to the stem-loop of the mRNA as it is transcribed, preventing polyadenylation[11] and recruiting factors, such as U7 snRNP, that trim the 3’-end of the pre-RNA to from the mature histone mRNA [12–16]; reviewed in[17]). Slbp remains bound to the histone mRNA throughout its lifetime and participates in its processing, translation and degradation.

Due to the stoichiometric nature of the relationship of Slbp with histone mRNAs, the levels of Slbp are believed to regulate the total level of histone mRNA that can accumulate in the cytoplasm [18]. Slbp therefore facilitates post-transcriptional regulation of histone mRNA levels and the incorporation of appropriate proportions of both replication and non-replication dependent histone variants into chromatin [17,18].

As well as being involved in regulating cell cycle progression, Slbp is itself regulated through the cell-cycle, with increasing levels accumulating during G1/S followed by rapid degradation at the end of S-phase [19,20]. Slbp levels/activity are regulated at the protein level by the ubiquitin proteolysis pathway, a process mediated by Cyclin A-CDK1 and CK2 dependent phosphorylation of two threonine residues in the TTP motif located within the amino terminus of SLBP [17–21].

Loss of Slbp in *C. elegans*, *Drosophila* and mouse leads to defects in cell-cycle dependent histone mRNA production and processing, resulting in the accumulation or depletion of unprocessed histone mRNA in the cytoplasm and a reduction in histone protein production [22–25]. Such changes in histone production cause problems with chromosome condensation and chromatin structure leading to cell cycle arrest and genomic instability [23,26]. Loss of maternal Slbp function in *C. elegans*, *Drosophila* and mice as well as Slbp2 in zebrafish leads to very early embryonic defects with embryogenesis arrested at mid-blastula transition (MBT; [22,24,27,28]. Transcriptomic analysis at MBT in *Drosophila* showed zygotic gene activation to be severely compromised [29]. Disruption of Slbp function at later stages has revealed some surprising phenotypes that suggest unexpectedly cell-type-specific roles: in *Drosophila*, most *slbp* homozygous null mutants perish at late pupal stage but some survive to adulthood and show female sterility [24]; loss of Cdl-1 (Slbp) in *C. elegans* results in defects in pharynx morphogenesis and body elongation[23]; and, in zebrafish *slpb* mutants survive until 5dpf and present defects in retinal development [10].

In this study, in addition to the initially observed retinal coloboma, we identify several other phenotypes affecting the eyes and central nervous system (CNS) in *slbp* mutants. These include deficits in axon guidance and pathway formation, particularly at midline commissures. Despite the relatively specific nature of the nervous system phenotypes, RNAseq analysis showed that gene expression in *ele* is very dysregulated. Many of the gene expression changes are consistent with cells failing to express differentiation-related genes while retaining expression of genes linked to proliferation. Consequently, the loss of *slbp* function likely affects modules of spatially and temporally regulated genes that mediate the transition from proliferation to differentiation. Indeed, we observe that whereas early born cells appear to differentiate, at later stages, cells both in the mesoderm and neuroectoderm fail to transition from proliferation to differentiation. This suggests that despite their specificity, some *ele* phenotypes are most likely a consequence of early born neurons differentiating within an environment that fails to mature appropriately.

## Results

### The *eisspalte* mutation is in *slbp*

To identify genes contributing to eye morphogenesis, we screened existing lines of fish carrying genetic mutations and noticed that homozygous *eisspalte^ty77e/ty77e^* (*ele*) mutant embryos frequently exhibited coloboma (Fig. 1A-B), a failure in closure of the optic/choroid fissure of the eye. The *ele* mutant was originally identified in a screen for mutations affecting brain morphology with the phenotype described as a dent in the midbrain-hindbrain boundary (MHB) [7]. In addition to this dent, slightly smaller eyes and a downward curve to the body axis were early morphological hallmarks of the *ele* phenotype, evident by 32hpf (not shown) and becoming more prominent by 2-3 days post-fertilisation (dpf; Fig 1A’). Failure of choroid fissure closure was evident by 2 dpf (Fig 1A’,B’). Two other phenotypes observed in the mutant at this late stage were heart oedema (Fig 1A) and abnormal otolith development, with otoliths smaller and fused together (not shown). Variations in penetrance and expressivity of *ele* phenotypes were observed in different genetic backgrounds. The *ele* phenotype was most severe in the TU strain such that although a downward curve to the body axis is present in TU, AB and WIK backgrounds, no obvious coloboma or MHB dent were observed when the mutation was crossed into the AB and WIK backgrounds.

**Figure 1.**
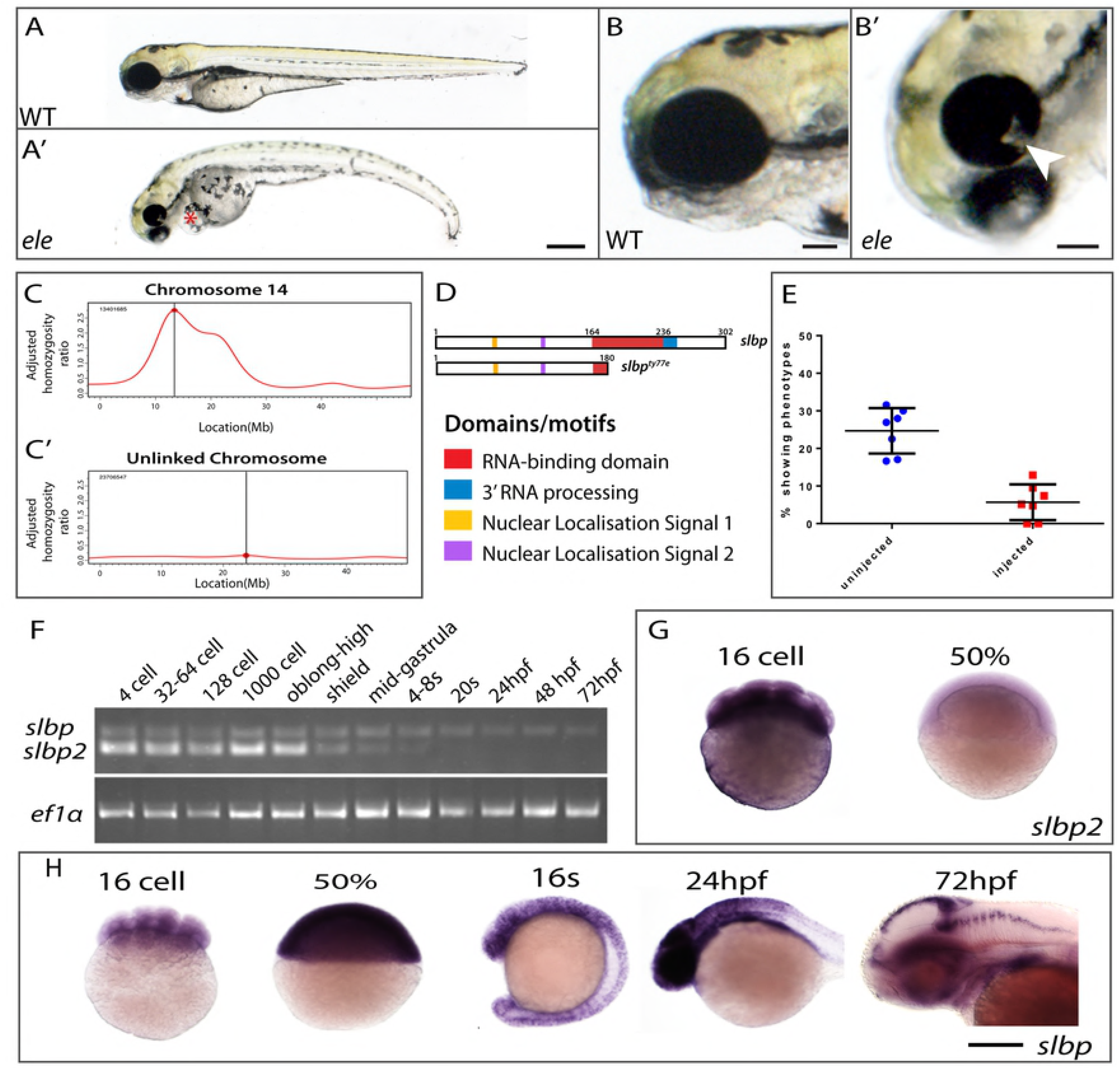
The *ele* mutation is in *slbp*. (A-A’) Morphology of 2dpf wildtype and *ele* mutant embryos (red asterisk indicates heart oedema). (B, B’) Heads of 2dpf wildtype (A) and *ele* (A*’*) mutant embryos showing the coloboma phenotype in the mutant (arrowhead in B’). (C-C’) RNAseq mapping plot of SNP homozygosity across Chromosome 14 and 15. (D) Domain organisation of wildtype and mutant Slbp predicted proteins. (E) Dot plots of percentages of embryos showing *ele* phenotypes in control clutches (blue) and in clutches injected with wildtype *slbp^TT-AA^*-RFP RNA (red). Each dot is the percentage from one of seven independent experiments. Thick black bars = standard deviation; fine black line = mean. (F) Reverse transcriptase polymerase chain reaction (RT-PCR) for *slbp* and *slbp2* at developmental stages indicated. (G, H) Whole mount RNA *in-situ* hybridization for *slbp* (H) and *slbp2* (I) at developmental stages indicated. Scale bars: (A, A’, H, I) 250μm; (B and B’) 100μm.

Bulked segregant analysis using SSLPs localised the *ele* mutation to chromosome14 between markers z4896 and z6847 (10.75Mb and 17.26 Mb respectively). This location (between 10 and 20Mb) was confirmed with an RNA-seq based mapping approach using a modification of the Cloudmap mapping pipeline on Galaxy (Fig 1C,C’; http://usegalaxy.org/cloudmap; [9]. RNA-seq data showed that the interval containing the *ele* mutation harboured 14 protein-coding (non-synonymous, splice, stop) variants of which only one with a stop codon (position 14.8Mb, located in the middle of the peak; Fig1C). The non-sense point mutation (C-to-A) is located in exon 5 of the *slbp* gene that encodes Stem-loop binding protein (Slbp), introducing a premature stop codon (Y180stop) within the 73 amino acid RNA binding domain (RBD) (Fig 1D). This mutation is predicted to lead to a truncation in Slbp at the amino-terminus of the RNA binding domain (RBD), generating a protein that lacks all the conserved residues required for RNA binding activity and histone pre-mRNA 3’UTR processing (Fig 1D). Supporting the possibility that the causative mutation is in *slbp*, embryos injected with a splice-site morpholino, targeting the exon3-to-intron3 donor site in *slbp*, phenocopied various aspects of the *ele* phenotype including the morphological dent caudal to the MHB and coloboma (data not shown). Furthermore, another published mutation (*slbp^rw440^*) in *slbp* has been shown to cause retinal defects [10].

To test whether reduction/loss of Slbp function causes all observed phenotypes, we injected synthetic RNA encoding wild-type RFP-tagged Slbp into embryos from a cross between *ele* heterozygous fish and assayed these embryos for rescue of phenotypes. Expression of Slbp-RFP fusion protein was confirmed by the presence of fluorescence at early gastrulation stages but no expression was detected by 26 hpf, suggesting that Slbp-RFP fusion protein may be degraded by this stage (data not shown). This is consistent with the notion that Slbp protein undergoes cell-cycle regulated cycles of synthesis and degradation [20,21,30]. To overcome these cycles of degradation, we therefore mutated two threonine residues within the TTP motif to alanines creating a construct encoding a degradation resistant Slbp^TT-AA^–RFP fusion protein[20].

Injected degradation-resistant *slbp^TT-AA^*-RFP RNA encodes a stably expressed nuclear localised protein that fully rescued the curve in the body axis, the MHB dent, commissural defects (described below) and the coloboma phenotypes over the first few days of development (data not shown). In control, non-injected embryos, 25.7% (*n*=100/389) showed an *ele* phenotype and the remaining embryos appeared normal consistent with full phenotypic penetrance in homozygous mutants. In the experimental group, injection of 150pg of degradation-resistant *slbp^TT-AA^*-RFP synthetic RNA reduced the number of embryos with an *ele* phenotype (as defined above) to 6% (Fig 1E; n=13/206 showed an *ele* phenotype; n=193/206 showed no phenotype). This rescue confirms that altered or absent Slbp function is the cause of the phenotypes we observe in mutants and for the remainder of this paper *ele* mutants will be referred to as *slbp^ty77e^* mutants.

### Only *slbp* and not *slbp2* is expressed in proliferative neural cells

To better understand how compromised Slbp function may lead to *slbp^ty77e^* phenotypes, we analysed the spatial and temporal expression of *slbp* and the paralogous *slbp2* gene. *slbp* and *slbp2* transcripts are maternally expressed and ubiquitously distributed in early cleavage stage embryos (Fig 1F-H and[10]). *slbp2* transcripts are undetectable by in situ hybridization from 50% epiboly stage but continue to be detected by RT-PCR until 4-8 somite stage suggesting gradual depletion of a maternal transcript pool (Fig 1F,G). In contrast, by 16s, increased levels of spatially restricted *slbp* transcripts were observed in the presumptive central nervous system and from 24hpf, high levels of expression start to become restricted to the proliferative zones of the brain, retina, fin buds and trunk (Fig. 1H; [10,31]). The expression of *slbp* is maintained in proliferative zones of the CNS at 2dpf (Fig 1H) where it overlaps with the expression of several replication dependent histone genes (e.g. *h1f0-H1*: Fig.S2A, B).

Both *slbp* and *slbp2* transcripts are maternally deposited suggesting that Slpb2 along with wildtype maternally derived Slbp may compensate for loss of functionality of zygotic Slbp during very early development. However as maternal RNA and protein is depleted and *slbp2* is not expressed after gastrula stages, this is likely to lead to the emergence of *slbp*^ty77e^ phenotypes in the nervous system during subsequent development.

### *slbp*^ty77e^ mutants have less neurons and show axonal defects

Coloboma phenotypes have been associated with retino-tectal pathfinding defects (eg. [32,33]) and indeed acetylated α-tubulin labelling of axons showed that retino-tectal projections are severely compromised in *slbp*^ty77e^ mutants. In wildtype animals, retinal ganglion cell (RGC) axons exit the eye via the choroid fissure at the optic nerve head, decussate at the optic chiasm and extend dorsally to innervate the contralateral optic tectum (Fig 2A,B). In *slbp*^ty77e^ mutants, the few RGC axons present often failed to exit the eye to form the optic nerve (Fig 2A’) and instead extended in aberrant locations within the retina itself (Fig 1C’ arrowhead; see also Imai et al, 2014). The optic tectal neuropil, formed by both RGC axons and tectal neuron dendrites, was severely depleted in *slbp*^ty77e^ mutants (Fig 2B’) suggesting that tectal neurons may also be depleted.

**Figure 2.**
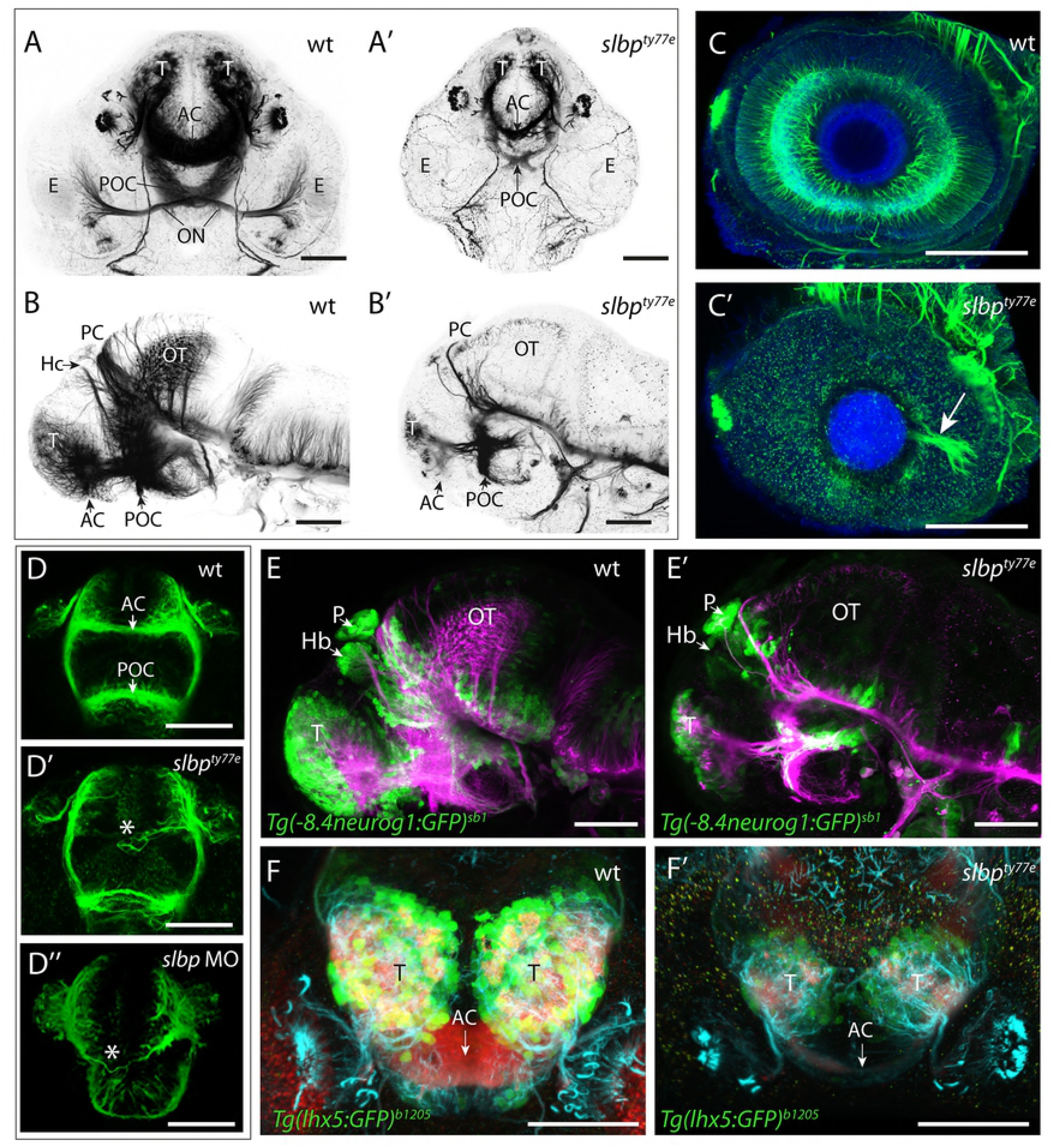
*slbp^ty77e^* mutants have less neurons and show axonal defects. (A-D”) Acetylated α-tubulin labelling of wildtype (A-D) and *ele* mutant (A’-D’’) embryo brains and eyes. Frontal (A-A’; D-D”) and lateral (B-B’) views of brains and lateral views of eyes (C,C’) of 3dpf (A-C’) and 30hpf (D-D’’) wildtype (A-D), *slbp^ty77e^* mutant (A’-D’) and morphant (D”) embryos. Arrowhead in C’ highlights the aberrant extension of RGC axons within the retina. The asterisks in D’-D” highlight aberrantly positioned axons near the midline commissural region in *slbp* mutant and morphant embryos. (E-E’) Lateral view with anterior to the left of 3dpf wildtype (E) and *slbp^ty77e^* mutant (E’) embryos showing expression of the *Tg(−8.4neurog1:GFP)^sb1^* transgene (green) labelled neurons and acetylated α-acetylated tubulin labelled axons/neurites (magenta). (F-F’) Frontal views of the telencephalon in 3dpf wildtype (F) and *slbp^ty77e^* mutant (F’) embryos showing expression of the Tg(lhx5:GFP)^*b1205*^transgene (green) labelled neurons, SV2 labelled neuropil (red) and acetylated α-acetylated tubulin labelled axons/neurites (cyan). Abbreviations: AC, anterior commissure; POC, post-optic commissure; ON, optic nerve; T, telencephalon; PC, posterior commissure; OT, optic tectum; Hb,habenula; E,eye; P,pineal. Scale bars: (A-C”, E, E’) 100μm (D-D”, F, F’) 50μm.

In addition to retino-tectal defects, several forebrain commissures, particularly those that form later in development [34,35], were reduced or absent in *slbp*^ty77e^ mutants/morphants. The brain underwent relatively normal morphogenesis and the anterior and post-optic commissures (AC and POC, respectively) formed but they were reduced in size and showed aberrant axons directed away from the main commissural pathway (Fig 2A’). In all genetic backgrounds analysed, the stria medullaris and tract of the habenula commissure did not form and consequently the habenula commissure (HC) was absent (Fig 2B’). In the TU background another dorsal commissure, the posterior commissure (PC) was present in *slbp*^ty77e^ mutants but was reduced to a thin bundle of axons crossing the midline, comparable to the tract in much younger wild type embryos. Additional axons were observed in the tract of the PC ventrally, but these did not extend dorsal-wards to the commissure (Fig 2B’). Posterior commissural defects were less severe in other genetic backgrounds in which more axons crossed the midline, though the commissure itself was less compact and axons were spread over a wider area (data not shown). These defects are consistent with commissure establishment and growth stalling after the first day of development.

Aberrant formation of the AC was the first discernable axonal defect we observed in *ele* mutants/morphants. In 30 hpf wildtype embryos, telencephalic axons have crossed the midline to form the AC (Fig 2D). In *slbp*^ty77e^ mutants/morphants this process was delayed and no AC was visible at 30hpf (Fig 2D’D”). The post optic commissure (POC) was present in *slbp*^ty77e^ mutants by this stage but was usually defasciculated (Fig 2D’). Correlating with these deficits, various genes encoding midline axon guidance molecules (including *sema3d, slit2, zic2.1* and *netrin1a*) showed misexpression in *slbp*^ty77e^ mutants (Fig S1). The severity of these commissural defects was variable in different genetic backgrounds and strongest in TU.

To assess whether the reduction in the extent of axonal labelling in *slbp*^ty77e^ mutants was correlated with a reduction in the numbers of neurons, we examined expression of the Tg(−8.4neurog1:GFP) transgene [36] which is present in many neurons throughout the anterior CNS and the Tg(lhx5*:GFP)^b1205^* transgene in dorsal telencephalic neurons[37]. Wildtype and mutant embryos labelled with antibodies to GFP and acetylated tubulin showed no overt difference in the extent and pattern of neurons in either Tg(−8.4neurog1:GFP)^sb1^ or Tg(lhx5*:GFP)^b1205^* backgrounds prior to 30hpf (data not shown). However at later stages, *slbp*^ty77e^ mutants (Fig 2E’, F’) had fewer Tg(−8.4neurog1:GFP)^sb1^ and Tg(lhx5*:GFP)^b1205^* positive neurons throughout the forebrain when compared to wildtype siblings (Fig 2E,F). A similar phenotype was seen in the retina in which the earliest born *ath5*:GFP^rw021^ expressing RGCs were observed in the ventro-nasal retina of mutants but later born neurons in the central retina were depleted (Fig 4A’) and subsequent waves of neurogenesis were delayed and less RGCs were present at later stages (data not shown). Later born retinal neurons were severely depleted in *slbp*^ty77e^ mutants with rods (not shown) and cone photoreceptors almost absent (Fig 4B’). Overall, neurons in *slbp*^ty77e^ mutant brains initially appeared relatively normal but after 30hpf, neuronal clusters failed to expand and late-born neurons were severely depleted/absent suggesting production of neurons may be arrested in the mutant.

To determine if the reduction in the number of neurons was due to an increase in programmed cell death, we performed TUNEL labelling on 30hpf embryos. An increase in TUNEL-labelled cells was observed in the tectum of *slbp*^ty77e^ mutants from 30 hpf (Fig 3A-A’). *slbp*^ty77e^ mutants showed no obvious cell death in the retina or forebrain at this stage and cell death in the lens comparable to wildtype siblings (Fig 3B-B’). We next asked if the axonal defects could be a consequence of the increased numbers of apoptotic cells in the brain. Blocking cell death from 16hpf using a caspase inhibitor did increase the level of acetylated tubulin labelling of neurites in the optic tectum and cerebellum (compare Fig 3C’ to 3C”) where apoptotic cells are prominent in *slbp*^ty77e^ mutants but did not rescue axonal defects. In such embryos, aberrantly located retinal axons were still present (Fig 3C’’) and habenular commissure and neuropil defects persisted. Overall the elaboration of axon tracts and neuropil did not recover to wildtype levels (Fig 3 C-C’’) in caspase inhibitor treated *slbp*^ty77e^mutants indicating that the axonal deficits seen in *slbp*^ty77e^mutants are not simply a consequence of cell death.

**Figure 3.**
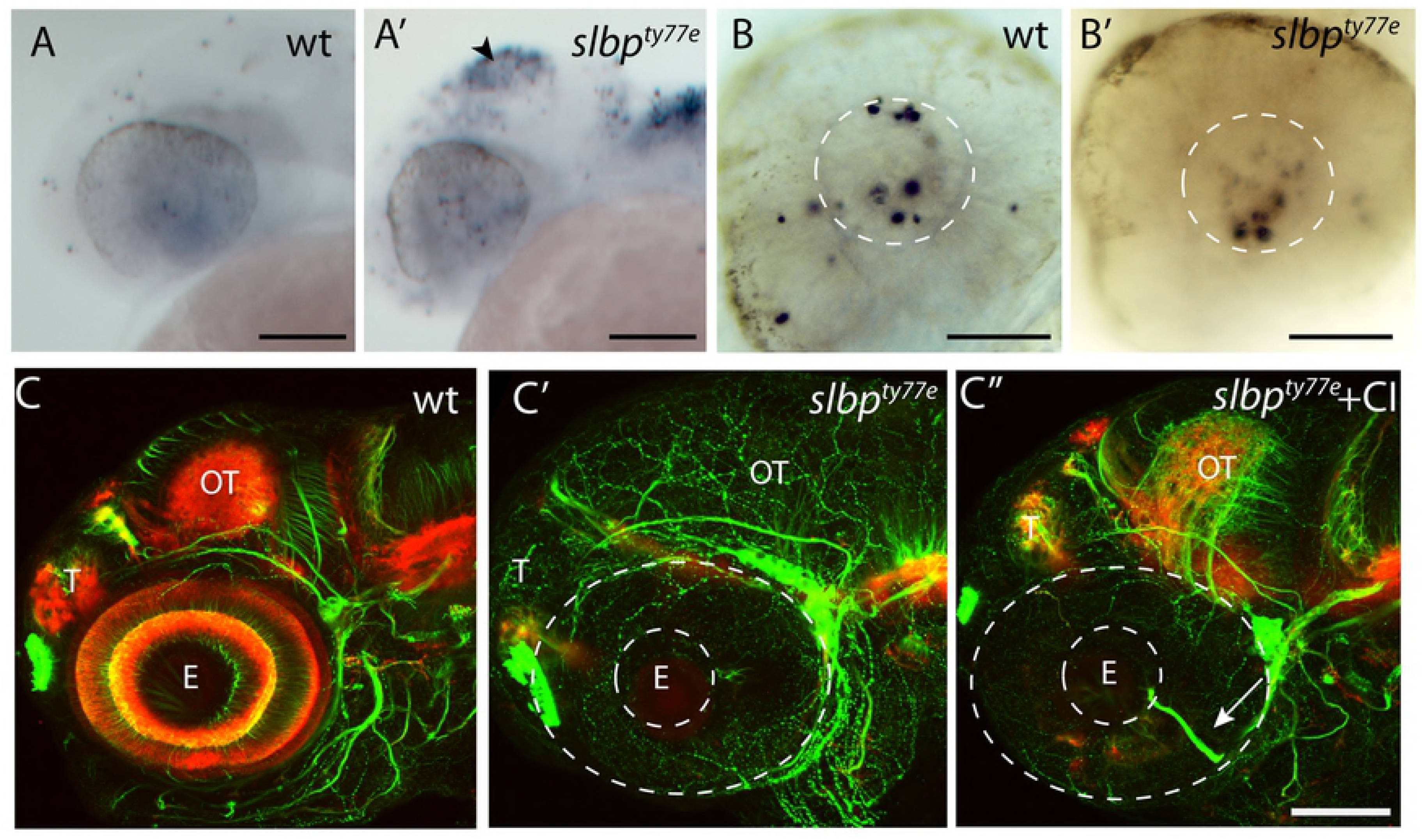
Suppressing cell death partially restores axonal/neuropil deficits in the tectum of *slbp^ty77e^* mutants. (A-B’) Lateral view of 26hpf wild type (A,B) and *slbp^ty77e^* (A’,B’) heads and eyes stained with TUNEL. Apoptosis is prominent in the tectum (arrowhead) and dorsal hindbrain of the mutant (A’) but not within the eye (B’). The white lines outline the lens within which there is apoptosis in both the wildtype and mutant eyes. (C-C”) Lateral view of 72hpf untreated wildtpe (C), untreated *slbp^ty77e^* (C’) and (C”) caspase inhibitor treated *slbp^ty77e^* heads labelled with anti-acetylated tubulin (axons, green) and anti-SV2 (neuropil, red). Arrow shows aberrant retinal ganglion axons in the mutant eye. Abbreviations: AC, anterior commisure; E, eye; OT, optic tectum; T, telencephalon. Scale bar: (A-A’,C-C”) 100μm; (B-B’) 50μm.

### Slbp is cell-autonomously required for differentiation and lamination of retinal neurons

To address whether the *slbp*^ty77e^ mutation compromises neuronal differentiation in a cell autonomous manner, we performed cell transplantation experiments in the eye in which patterns of neurogenesis are very stereotyped and differentiated neuronal types are easy to visualise.

GFP-expressing wildtype cells transplanted into wildtype retinas formed clonal columns of retinal neurons, with cells occupying all layers of the retina (n=30, Fig 4C,D). In contrast, *slbp*^ty77e^ mutant cells transplanted into wildtype retinas lacked neuronal morphologies and appeared clumped instead of being distributed throughout the three layers of the retina (n=7 clones in 4 retinas; Fig 4C’,D’). Notably, *slbp*^ty77e^ mutant cells rarely, if ever, contributed to the outer neural retina in which many later born neurons reside. Moreover, retinal lamination, visualised with β-catenin antibody, was absent within, and adjacent to, the *slbp*^ty77e^ mutant clones (Fig 4D’). These experiments show that Slbp is required cell autonomously for differentiation and lamination of retinal neurons and that clones of *slbp* mutant cells can non-autonomously disrupt organisation of adjacent wild-type retina.

**Figure 4.**
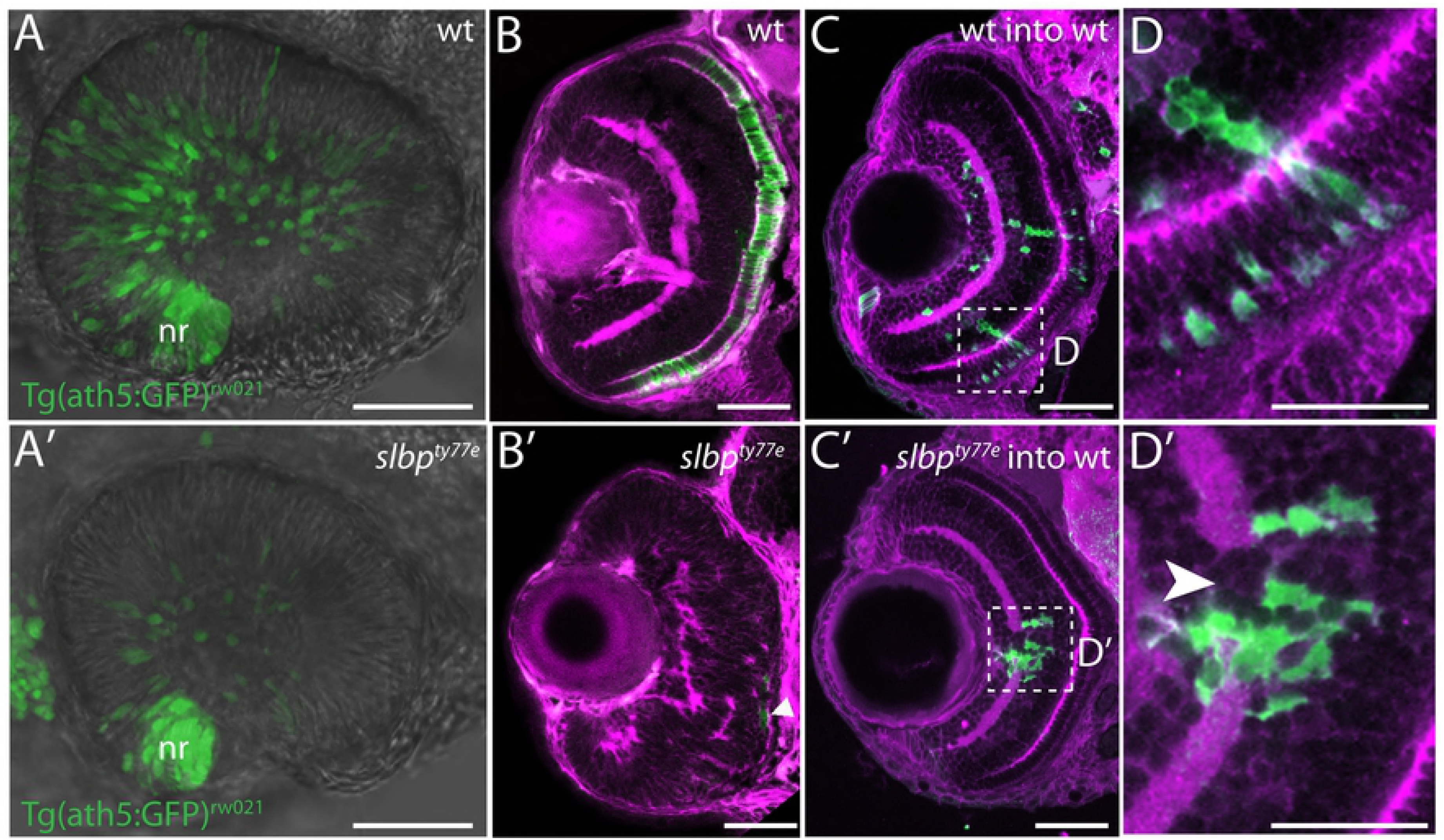
Slbp is required cell autonomously for retinal neuron differentiation. (A, A’) Images from live timelapse recordings of *Tg(atoh7:GFP*)^rw021^ GFP-transgene expression in retinal neurons in control (A) and *slbp^ty77e^* (A’) eyes at 30hpf. (B-B’) Frontal sections of 3dpf wildtype (B) and *slbp^ty77e^* (B’) retinas showing anti-γ-tubulin labelled neurites/neuropil (magenta) and zpr1-expressing cone photoreceptors (green). The arrowhead points to a few remaining zpr1+ cone cells in the *slbp^ty77e^* mutant eye. (C-C’) Frontal sections of 3dpf wildtype (wt) host retinas containing transplanted wildtype (C) or *slbp^ty77e^* mutant (C’) GFP-labelled cells (green). (D-D’) High magnification images of transplanted cells in C-C’. Note the break in the plexiform layer in the retina (arrowhead) in the vicinity of the *slbp^ty77e^* mutant cells. Abbreviations: nr, nasal retina. Scale bars: (A-C’) 50μm; (D-D’) 25μm.

### *slbp*^ty77e^ mutant cells fail to transition from proliferation to differentiation

Slbp regulates histone mRNA metabolism and levels of Slbp protein are tightly cell cycle regulated [18,20,22,24,27]. To examine the effect of loss of Slbp function on cell cycling we first utilized flow cytometry to profile cell phasing in dissociated cells from 2dpf wildtype and *slbp*^ty77e^ mutant embryos. The percentage of *slbp*^ty77e^ mutant cells (47.4%) in S-phase was double that in wildtype (23.6%). Similarly, 10.4% of *slbp*^ty77e^ mutant cells were in G2/M phase compared with 4.1% of wildtype cells. Conversely, fewer *slbp*^ty77e^ mutant (41%) than wildtype (71.7%) cells with G1 DNA content were observed (Fig 5A).

**Figure 5.**
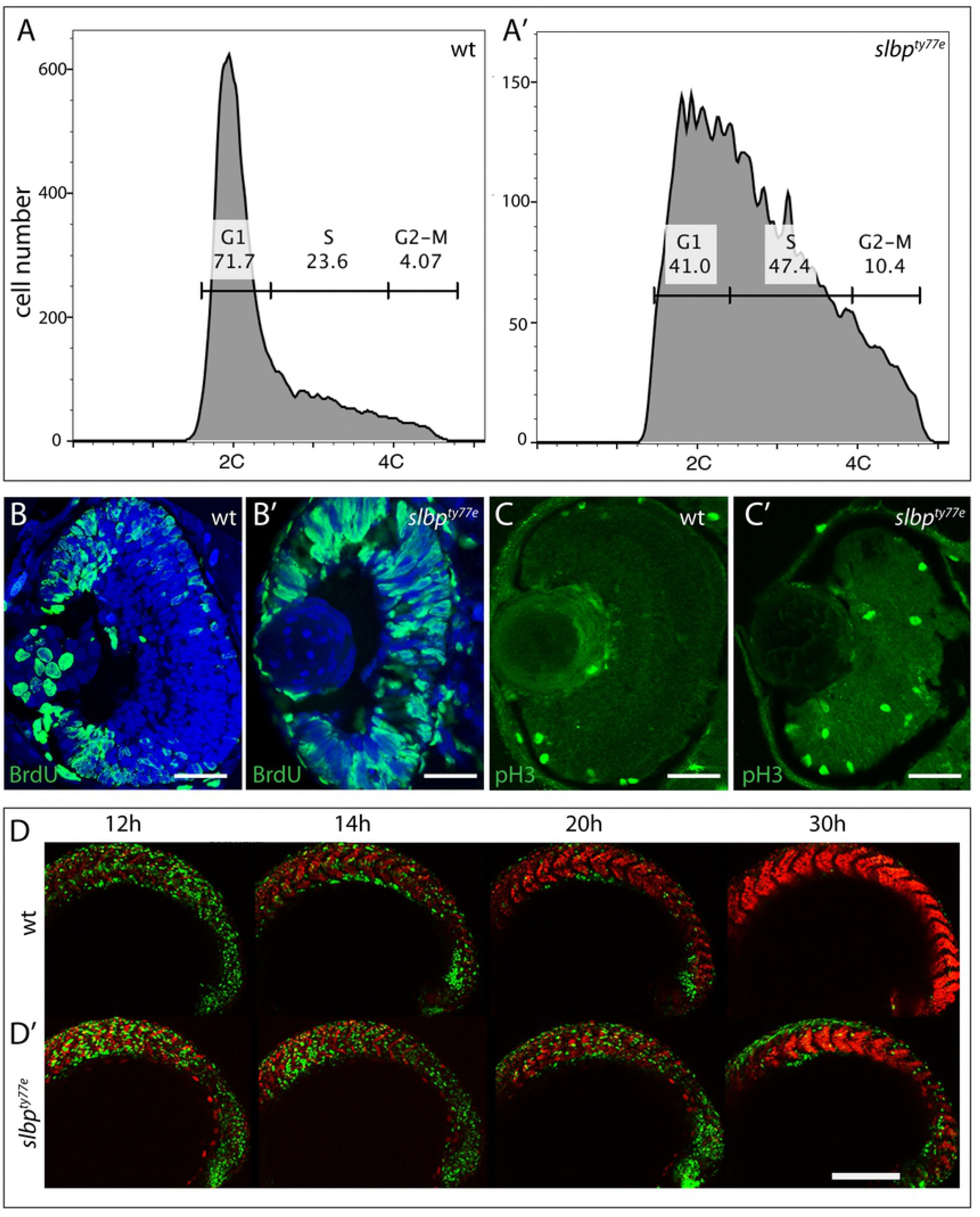
*slbp^ty77e^* mutant cells fail to transition from proliferation to differentiation. (A,A’) DNA content of 48 hpf wildtype and *slbp^ty77e^* embryos as assessed by flow cytometry. (B,B’, C,C’) Transverse sections through 48hpf wildtype (B,C) and *slbp^ty77e^* mutant (B’,C’) eyes immunostained with anti-BrdU (B,B’) and anti-pH3 antibodies (C,C’). (D,D’) Series of projections of confocal images extracted from live time-lapse movies of *tg(EF1α:mAG-zGem(1/100))^rw0410h^*^;^ *tg(EF1α: mKO2-zCdt(1/90))^rw0405b^* transgenic wildtype (D) and *slbp^ty77e^* mutant (D’) embryos showing cell cycle progression in cells of the forming somites. Green cells are proliferative (S, G2, M) whereas red cells are differentiating somite cells. Scale bar: (B-C’) 50μm; (D-D’) 300μm.

To assess if proliferative cells showed abnormal spatial distributions in *slbp*^ty77e^ mutants, we assessed BrdU incorporation (which labels cells in S phase) and PH3 labelling (which recognises mitotic cells). In wildtype 56hpf embryos, S and M-phase proliferating retinal cells were confined to the ciliary marginal zone whereas many BrdU and PH3 positive cells were located in the central retina of *slbp*^ty77e^ mutants (Fig 5B-C’) as has also been shown in *slbp^rw440^* mutants[10]. Furthermore, expression analyses that showed that cyclins representative of all stages of the cell cycle *cyclin D1*, *cyclin E2*, *cyclin A, cyclin B* were all upregulated in *slbp*^ty77e^ mutants and were expressed in areas of the brain and eye that are largely post mitotic in wild-type embryos (Fig 6, S2 and data not shown). These results suggest that many neural cells remain proliferative in *slbp*^ty77e^ mutants and fail to transition to generating post-mitotic neurons.

**Figure 6.**
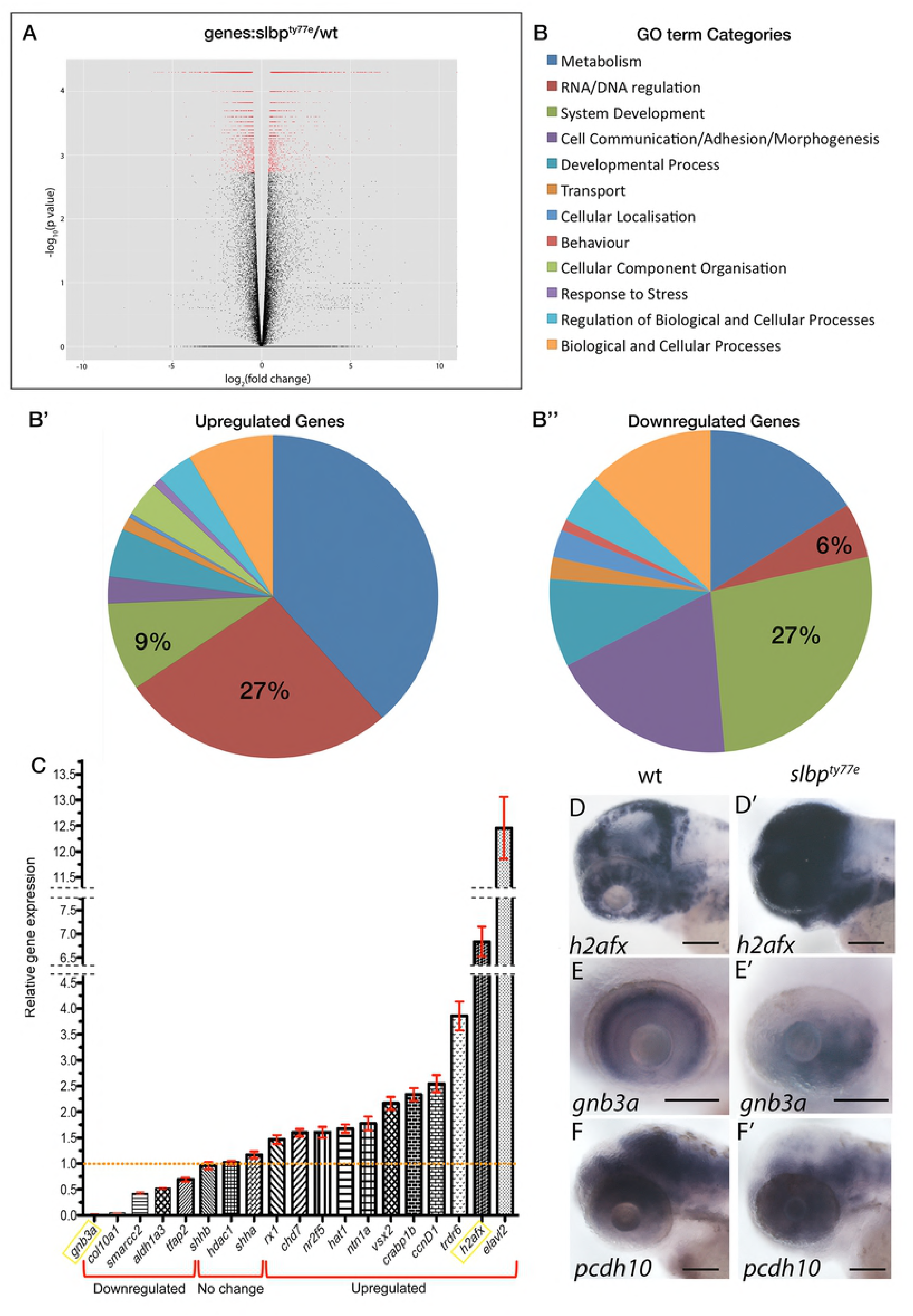
RNAseq analysis reveals that loss of Slbp function induces large-scale changes in gene expression levels consistent with many cells failing to transition to differentiation. (A) Volcano plot displaying differential expressed genes between wildtype and *slbp^ty77e^* embryos The red dots on the right represent the significant up regulated expressed transcripts (p < 0.01, false discovery rate (FDR) q < 0.01); the red dots on the left represent the transcripts with expression significantly down regulated (p < 0.01, FDR q < 0.01). Non-significant genes (q > 0.01) are represented by a black dot. (B) GO term categories for enriched genes. (B’, B”) Pie charts showing percentages of GO terms relating to each category for up (B) and down (B’)-regulated genes in *slbp^ty77e^* mutants compared to wildtype. C) Graph showing real time PCR quantification of expression changes for genes selected from the RNAseq dataset. Samples were normalized to *β*-actin and wildtype values for each gene were set to 1. Fold changes in mutants were plotted relative to this value. (D-F’) Lateral views of 3dpf wildtype (D-F) and *slbp^ty77e^* mutant (D’-F’) heads/eyes showing expression of *h2afx* (D, D’); *gnb3a* (E,E’) and *pcdh10* (F,F’). Note that expression changes for *h2afx* and *gnb3a* (C) are consistent with qPCR data (yellow box in C). Scale bars: 100μm.

To define the onset of cell cycle defects in *slbp*^ty77e^ mutants, the *Tg(EF1α:mAG-zGem(1/100))^rw0410h^* transgene [38] that visualizes cells in S, G2 and M phases (Fucci green) and *Tg (EF1α: mKO2-zCdt(1/90))^rw0405b^* transgene that highlights cells in G1 phase (Fucci orange) were crossed into fish carrying the *slbp*^ty77e^ mutation and embryos analysed to detect the ratio of proliferating and non-proliferating cells in live embryos. These transgenes were expressed at low/negligible levels in the nervous system and so our analysis focussed on the transition from proliferation to differentiation in the mesodermal somites.

Timelapse imaging of the developing somites showed that from about 14 hpf, most cells in anterior somites (the earliest forming) of wildtype embryos were post mitotic (red) whereas in *slbp*^ty77e^ embryos, the majority of cells were still expressing transgenes normally restricted to proliferating cells (Fig 5D-D’). Some somitic *slbp*^ty77e^ cells maintained expression of S/G2/M phase transgenes through later developmental stages (30/32hpf; Fig 5D-D’). Consequently, although *slbp*^ty77e^ mutants only show an overt morphological phenotype from around 30-32hpf, defects in the transition from proliferation to differentiation are already present from as early as 12/14 hpf. These results are consistent with the observations above and suggest that both in mesodermal and ectodermal tissues, *slbp*^ty77e^ mutant cells are compromised in their ability to effectively transition from proliferation to differentiation.

### RNAseq analysis of *slbp*^ty77e^ mutants reveals mis-regulation of histone and chromatin remodelling genes and loss of expression of genes indicative of differentiation

RNAseq analysis of *slbp*^ty77e^ mutant and wild-type embryos at 52hpf showed that gene expression in *slbp*^ty77e^ is strongly dysregulated in mutants with 2158 genes significantly upregulated (Table S2) and 2607(Table S3) genes downregulated based on a q value ≤ 0.01 (Fig 6A). Consistent with the role of Slbp in histone RNA processing [18], we found that histone transcripts were highly enriched in *slbp*^ty77e^ mutants including both canonical and non-canonical histone subunits (49 of the top 100 highest fold over-represented transcripts are from histone subunit encoding genes).

To determine whether the expression of certain groups of genes is particularly dysregulated in *slbp*^ty77e^ mutants we used the AmiGO2 tool (The Gene Ontology Consortium) [39] to perform a GO term enrichment analysis for “Biological Process” on genes showing a significant change in expression (q value ≤ 0.01) in the RNAseq data. This enrichment analysis was performed for both up (Fig 6B’, Table S2) and down (Fig 6B”, Table S3)-regulated genes. GO terms significantly enriched (p ≤ 0.05) in *slbp*^ty77e^ sequence datasets were compared to GO terms assigned to 25,800 protein-coding *danio rerio* genes and were manually grouped into 14 categories (Fig 6B; Table S2 & S3). GO terms relating to DNA/RNA regulation (many of which relate to chromatin regulation and cell cycle) were enriched, indeed accounting for 27% of enriched GO terms in the up-regulated gene list and 6% for down-regulated genes. GO terms relating to nervous system development (GO:0007399), neurogenesis (GO:0022008), eye development (GO:0001654) and axogenesis (GO:0007409) (grouped under GO term category; “system development”) were also highly represented in both upregulated (9% of all enriched GO terms) and downregulated (27% of all enriched GO terms) gene datasets.

Further interrogation of the lists above for genes expressed with a log fold change of greater than −2 revealed downregulation of many genes normally expressed in differentiated neurons (Table S4). For instance *crx* (Cone-Rod Homeobox Protein); *syt5b* (synaptotagmin Vb), *neurod6a* (neuronal differentiation 6a), *olig1* (oligodendrocyte transcription factor 1); *slc6a1b* (solute carrier family 6 (neurotransmitter transporter, member 1b); *nr4a2a* (nuclear receptor subfamily 4, group A, member 2a); *gria1a* (glutamate receptor, ionotropic, AMPA1a); *slitrk5a* (SLIT and NTRK-like family, member 5a); *gad2*(glutamate decarboxylase 2); *grm2a* (glutamate receptor, metabolic 2a) were among the downregulated genes normally expressed in mature neurons. In contrast among those genes comparably upregulated were many linked to cell cycle and proliferation (Table S4 and Fig S2). One of the most upregulated genes in *slbp*^ty77e^ mutants was *elavl2*, which encodes another RNA binding protein [40] that is expressed as progenitors transition to post-mitotic neurons ([41]; Reviewed in[42]) while expression of both *elavl3* and *elavl4*, expressed in mature neurons, was downregulated. These results are consistent with many *slbp*^ty77e^ mutant cells failing to differentiate and maintaining expression of genes characteristic of proliferative progenitor cells.

To validate and further interrogate the RNAseq data, we assessed expression of several genes related to histone function or eye/nervous system development by in situ hybridization and/or quantitative RT-PCR. qPCR of 19 selected genes showed comparable upregulation or downregulation of expression as observed in the RNAseq dataset (Fig 6C). In situ hybridisation analysis of 10 genes from the RNAseq dataset (of which *gnb3a*, c*yclinD1*, *netrin1a* and *h2afx* were also analysed by qPCR) showed changes in expression that confirmed RNAseq data (Fig 6D-F’ and Fig S1, S2). For instance, instead of being restricted to expected sites of neural cell proliferation, histone *h2afx* mRNA was expressed very broadly throughout much of the brain whereas the Protocadherin encoding gene *pcdh10*, which is expressed in neurons and regulates axon guidance [43], was widely downregulated. Together, RNAseq data confirmed by qPCR and ISH showed widespread developmental transcriptional dysregulation in *slbp*^ty77e^ mutants, with many changes consistent with a failure of many cells to transition to differentiation.

Altogether, these results support that the axonal and retinal phenotypes observed in *slbp*^ty77e^ mutants are potentially due to mis-regulation of modules of regulatory genes important for the transition from proliferation to differentiation and subsequently for specific features of nervous system development such as axon guidance and retinal morphogenesis.

## Discussion

In this study we describe the cloning of the *slbp*^ty77e^ mutation and characterization of nervous system phenotypes in *slbp*^ty77e^ mutants. We show that although early born neurons are present and elaborate axons in *slbp*^ty77e^ mutants, later born neurons are severely depleted and consequently late forming commissures are absent and early tracts and commissures fail to grow. These results are consistent with the observation that many proliferative neural cells fail to transition to differentiation and consequently there are major alterations in the spatial and temporal distribution of proliferative versus differentiated cells in the developing nervous system. One result of this phenotype is that early born neurons differentiate in a very abnormal environment and this no doubt contributes to the axon guidance and other defects present in mutants. Such phenotypes may be a consequence of the role of Slbp in regulating histones that modulate chromatin structure thereby influencing expression of modules of developmental regulatory genes.

### A failure in transitioning from proliferation to differentiation may underlie most *slbp* mutant nervous system phenotypes

While *slbp* mutants are not viable in mammals [22,44,45], the survival of fish *slbp* mutants through embryogenesis allowed us to study relatively late nervous system phenotypes. The late appearance of phenotypes in fish *slbp* mutants is likely due to the presence of both maternally provided *slbp* transcripts and the early expression of the paralogous *slbp2* gene through gastrulation stages. We assume that nervous system phenotypes emerge after the gradual depletion of these pools of Slbp/Slbp2 protein.

A consistent feature of our phenotypic analyses of *slbp* mutants is the depletion of neurons and continued presence of cells that fail to transition from proliferation to differentiation. For instance, within *slbp* mutant eyes, the earliest born retinal neurons in the ventro-nasal retina appear as normal but later born neurons in the central retina are severely depleted; in contrast BrdU-incorporating cells and mitotic figures remain present within the central retina, long after they are largely absent in wild-type eyes. This phenotype is consistent with other studies/contexts in which loss of Slbp has been linked to cell cycle progression and differentiation deficits [10,46,47].

The presence of early born neurons and early axon tracts and absence of later neurons/tracts is perhaps most simply explained if sufficient maternal Slbp/Slbp2 protein is retained in the precursors of these early-born neurons to enable them to exit the cell cycle. If this is indeed correct, then small differences in the levels/perdurance of maternal Slbp/Slbp2 protein may contribute to the variation in phenotypic severity seen in different backgrounds.

Although difficult to determine with any certainty, it is also possible that late phenotypes such as a failure in the closure of the choroid fissure and commissural axon guidance phenotypes could be a secondary consequence of the failure of many neural cells to differentiate. For instance, choroid fissure closure is dependent on appropriately timed expression of genes in neural retina and retinal pigment epithelium [33,48]; and GG and SW, unpublished data); depigmentation of the ventral eye is associated with coloboma in *slbp* mutants suggesting a failure in differentiation of the retinal pigment epithelium cells may contribute to the coloboma. Similarly, the altered environment along the pathways through which axons from early born neurons extend almost certainly contributes to the observed axon guidance phenotypes. The early axon pathways in the brain are mostly established along boundary regions within the neuroepithelium, many of which are sites of neuronal differentiation[49]. The depletion of neurons in mutants coupled with the widespread misregulation of genes encoding axon guidance proteins expressed in neuroepithelial cells presents the extending axons with a very abnormal environment, no doubt contributing to their projection errors.

### Slbp regulates expression levels of numerous histone genes and other genes affecting chromatin

In *slbp*^ty77e^ mutants, the *slbp* gene encodes a predicted protein that is truncated at the amino-terminus of the RNA-binding domain and lacks all the conserved residues required for RNA binding activity and histone pre-mRNA 3’UTR processing. Therefore, *slbp*^ty77e^ mutant Slbp is most likely devoid of all RNA binding activity. One consequence of this is a likely shortage of histone proteins during S phase, which would lead to aberrant chromatin structure. Indeed, RNAseq results showed genes encoding or associated with histone proteins are highly represented suggesting that the aberrant translational regulation of Slbp-dependent histones leads to profound changes in histone gene transcription. The upregulation of histone RNA expression could be due to the production of unprocessed and aberrant polyadenylated histone mRNAs that are more stable than corresponding wildtype RNAs as previously shown for histone H3 and H4 in *Drosophila* Slbp mutants [24] and for all replication-dependent histones in human cells [24]. This is in apparent contrast to recent work, again in *Drosophila*, showing that histone mRNA levels can be dramatically decreased in Slbp mutants [29]. Additionally selective downregulation of particular histones has been reported in mouse and *slbp2* zebrafish mutants [22,28].

### Degradation of SIbp at the end of S phase may not be essential for its function

In mammalian cells, Slbp levels are regulated in a cell-cycle dependent manner through a highly conserved phosphorylation motif (TTP) that targets Slbp for ubiquitin mediated degradation by the proteosome at the end of S-phase [20]. This motif is present in Slbp, (but surprisingly absent from maternally deposited Slbp2, [20]), suggesting that zebrafish Slbp has the potential to be regulated in an identical manner. Indeed, exogenously expressed wildtype Slbp is rapidly turned over (and consequently cannot rescue *slbp*^ty77e^ mutant phenotypes). However, in contrast, expression of a degradation resistant Slbp (*slbp1^TT-AA^*) very effectively rescues *slbp*^ty77e^ phenotypes. The simplest explanation of this result is that degradation of Slbp at the end of S phase is not required for histone mRNA regulation. Moreover, no overt phenotype was observed in wildtype embryos overexpressing *slbp1^TT-AA^* suggesting that embryos can tolerate excess Slbp throughout many cell cycles.

### Disrupted chromatin regulation underlies many congenital abnormalities of visual system and brain development

If the role of Slbp in chromatin regulation underlies some of the more intriguing phenotypes in *slbp*^ty77e^ mutants, then it would be consistent with an ever-increasing list of chromatin regulators being linked to human congenital abnormalities of eye and brain development [50–56]). For instance, loss of function of chromodomain helicase DNA binding protein 7 (CHD7, [57]; reviewed in [58] is the cause of CHARGE syndrome, a rare genetic syndrome that shares phenotypic characteristics with *slbp*^ty77e^ mutants. For instance, patients show congenital abnormalities in the visual system and brain including coloboma, cranial nerve deficits and intellectual disability (reviewed in [58]). Chd7 is also required for proper extension, pruning, guidance and extension of axons in the developing central nervous system of the fly [50,59], suggesting that such defects could contribute to the neurological symptoms in human patients.

Coloboma, small eyes, ear and neurogenesis defects are also observed when Hdac1 function is compromised [60–62]. Hdacs (Histone deacetylases) are among the most critical histone-modifying enzymes and their loss of function results in chromatin de-compaction and transcriptional perturbation (reviewed in [63]. Various studies have linked Hdac (and indeed Slbp) function with specific developmental genes and pathways such as Fgf [62,64], Notch [10,62] and Wnt [4,65,66] but as we show, it is possible to observe quite specific phenotypes even in contexts when there is massive dysregulation of gene expression. Consequently, when chromatin regulators are implicated in developmental events, transcriptomic studies provide a valuable overview of the expression landscape within which specific phenotypes may arise.

## Materials and Methods

### Zebrafish lines and genotyping

*AB*, *TU, WIK* and *EKWILL* wild-type, *eisspalte* (*ele/slbp^ty77e^*), *Tg(−8.4neurog1:GFP)^sb1^*[36]*, Tg(lhx5:GFP)^b1205^*[37], *Tg(atoh7:GFP)^rw021^* [67], *Tg(EF1α:mAG-zGem(1/100))^rw0410h^* and *Tg4(Xla.Eef1a1:mKOFP2-cdt1)^rw0405^b* [38] zebrafish (*Danio rerio*) lines were bred and maintained according to standard procedures [68]. To prevent pigment formation, 0.003% phenylthiourea (PTU, Sigma) was added to the fish water between 20 and 24hpf. For live timelapse imaging of Fucci lines embryos were anaesthetised in 0.2% tricaine methanesulfonate (MS222, Sigma) in fish water.

Genotyping of the *ele^ty77e^* mutation was performed following PCR analysis of genomic DNA using primers JH-641 (forward 5′-CTCATCAGAAGACAGAAGCAGATCAA**C**TA −3′) and JH-209 (reverse 5′-TTGCCCACCCCTGTTCTA-3′) followed by *Dde*I restriction digestion of the PCR products to generate 445 bp and 418 bp fragments for the wildtype and mutant alleles respectively. The bold C nucleotide is changed within the primer to create the *Dde*I restriction site in the *ele^ty77e^* mutant allele. Latterly, a KASP assay (KASP, LGC genomics; ID 1234567890), performed according to the manufacturer instructions was also used for genotyping embryos.

### SNP-mapping

*ele* heterozygotes in a TU background were outcrossed to WIK strain for bulk segregant linkage analysis [69] and to EKWILL for subsequent mapping. Simple sequence length polymorphisms (SSLPs) were used to establish low resolution linkage[70]. Single nucleotide polymorphisms (SNPs) were identified by sequence analysis of PCR products derived from heterozygote F1 *ele*/EKWILL fin-clip derived DNA for distantly located markers and by comparison to homozygous F2 EKWILL/EKWILL wildtype and *ele* mutant embryos derived DNA sequences for closely located markers. PCR products harbouring SNPs that gave rise to restriction fragment length polymorphisms (RFLPs) were digested with appropriate restriction enzymes and resolved by 2-4% agarose gel electrophoresis. If no restriction site was present, dCAPS Finder 2.0 software [71] was used to design primers that generate a restriction site polymorphism for analysis in the same manner. Sequence data was analysed using Lasergene Navigator software.

### Mapping-by-sequencing and RNAseq

We also used an RNAseq approach to map the *ele* mutation, identify causal variants and reveal gene expression differences in *ele* mutants. To obtain embryos at the same developmental stage, *ele^ty77^* heterozygotes were kept apart in breeding tanks and embryos collected 30 minutes after divider removal. Mutant and sibling embryos were sorted by phenotype at 2 dpf. For RNAseq analysis, we performed 2 biological replicates and 1 experimental replicate. Total RNA was isolated from 30 embryos using 500μl of Trizol followed by homogenization with a G30 syringe and standard chloroform extraction and ethanol precipitation. RNA integrity was validated by RQI > 9.6, where 0 corresponds to fully degraded RNA and 10 corresponds to intact RNA (Experion RNA HighSens Analysis, BIORAD).

RNAseq analysis was performed on an in-house Galaxy server using the Tuxedo pipeline [72]. Briefly, reads from both mutants and siblings were mapped to the zebrafish Zv9.65 genome using TopHat2, assembled into a parsimonious list of transcripts using Cufflinks and a merged transcript dataset from all the Cufflinks transcripts was created using Cuffmerge [72]. Differential expression analysis was performed on the BAM files from all three biological replicates and the merged transcript dataset using Cuffdiff (Table S1). Differential expression between mutant and sibling samples was only counted as significant if q < 0.01. GO term enrichment analysis for “Biological Process” on all of the genes showing a significant change in expression (q value ≤ 0.01) in our RNAseq data was performed using the AmiGO2 tool (The Gene Ontology Consortium) [39]. GO terms significantly enriched (p ≤ 0.05) in *ele* compared to a background list of 25,800 protein-coding *Danio rerio* genes were manually grouped into 14 categories (Table S2 & S3).

Mapping-by-sequencing was performed in parallel on the same in-house Galaxy server using a modified version of the Cloudmap variant discovery mapping (VDM) platform[8,9] to process RNAseq rather than whole-genome sequencing data. Instead of plotting individual allele frequencies of the variants identified in the mutant sample as per the VDM pipeline, the kernel density of homozygous/heterozygous SNPs was plotted along each chromosome. To narrow the list of causal variants in the mutant sample we subtracted homozygous variants identified in the sibling sample, as well as a list of common wildtype variants complied by combining variants identified through our own sequencing the *ekwill* strain plus a list compiled from previously published data [73,74].

### Microinjection

An *slbp* antisense morpholino oligonucleotide was designed to target the third coding exon-intron boundary and synthesised by Gene Tools LLC. *slbp* morpholino (5′-ATTCAAGAGAGGCAACTGACCGATG-3′) was injected at 2ng and to assess effectiveness of the morpholino to block splicing, RT-PCR analysis was carried out using primers JH711 (forward: 5′-CAAAGGAGCTTCAAGGTGGT-3′) and JH712 (reverse: 5′-AGGGAAATCACTCGCAAGAA-3′).

To generate a degradation resistant wildtype *slbp* expression construct for rescue analysis, (T92A) and (T93A) mutations were created using a PCR-based mutagenesis method [75]. The resulting fragment was cloned into CS2+RFP to generate CS2+RFP-Slbp^TT-AA^. Capped mRNA was prepared using the mMachine RNA Synthesis Kit (Ambion) according to the manufacturer’s instructions. One-cell-stage embryos resulting from *ele^ty77e^* heterozygous in-crosses were injected with 50pg of CS2+RFP-Slbp^TT-AA^ mRNA.

### DNA content analysis

To obtain single cell suspensions, between 40 and 50 anesthetized mutant and wild-type 48hpf embryos were incubated for 20 minutes on a shaker in 0.25% trypsin in L15 tissue culture media (Sigma). Repeated trituration using fire-polished glass pipettes was performed. Cell suspensions were cleaned with a mesh and re-suspended in PBS. Cells were fixed in 70% EtOH and stored at 4°C for several days. Cells were re-suspended in 100 microliters of propidium iodide solution (in 4 mM citrate buffer, pH 6.5, containing 0.1 mg/ml propidium iodide (Sigma), 200 μg/ml RNase, and 0.1% Triton X-100) and stored at 4°C until analysis. Data acquisition was performed by using a Becton Dickinson FACS-Calibur machine and analyzed by using the FlowJO programme.

### Histology

Whole-mount immunolabelling procedures were performed as previously described [34,76]. For antibody staining of cryosections, embryos were first protected by sequential incubation in 15% then 30% sucrose in phosphate-buffered saline supplemented with 0.5% Triton X-100 (PBST) for 12-16 hours at 4°C, embedded in OCT, stored at –80°C, and sectioned at 16-20μm using a Leica cryostat. γ-tubulin (Sigma; 1:200); zn5 [Zebrafish International Resource Center (ZIRC); 1:250]; GFP (AMS Biotechnology TP401; 1:1000); anti-acetylated tubulin (IgG2b, Sigma; 1:500); anti-SV2 (IgG1, DSHB; 1:500); BrdU (Roche; 1:300); PH3 (Upstate Biochemical; 1:500); zpr-1 (ZIRC, 1:50) antibodies were used.

Antisense mRNA probes for whole-mount in situ hybridisation were synthesised using RNA polymerases (Promega) and digoxigenin labelled nucleotides (Roche) following manufacturer’s instructions. Whole-mount in situ hybridisations were performed essentially as previously described [33]. TUNEL labelling to detect apoptosis was performed using the ApopTag Kit (Chemicon International). In ordered to block apoptosis, 24hpf embryos were treated with 300 μM of caspase inhibitor (Z-VAD-FMK, Sigma).

### Cell proliferation assays

BrdU (Sigma) incorporation was performed as previously described [77]. Briefly, de-chorionated embryos were incubated in 10mg/ml BrdU/15%DMSO in E3 on ice for 20 minutes, washed in warm E3 at 28 degrees for 20 minutes prior to fixation with 4% paraformaldehyde.

### Cell transplantations

Embryos from *ele^ty77^e* heterozygote incrosses were injected with GFP mRNA (40-50 pg per embryo) at the one-cell stage. Thirty to 40 GFP+ cells were transplanted from the apical region of mid-blastula donor embryos into early-gastrula-staged hosts in the region fated to become the eye [78,79]. Donor embryos were either genotyped or allowed to grow until 3 dpf to distinguish mutants from siblings. Host embryos were fixed at the stages indicated in the results, genotyped if necessary, then prepared for cryo-sectioning and antibody staining.

### Imaging and data processing

Embryos subjected to whole-mount in situ hybridisation were cleared in serial incubations of glycerol (25, 50, 75 and 95%), the eyes and brains dissected and placed in a drop of glycerol, cover-slipped, and imaged with a 40X (0.8 NA) water-immersion lens using a Nikon E1000 microscope connected to a digital camera (Jenoptik) operated by Openlab (Improvision) software.

Cryosections were examined by confocal fluorescence microscopy (Leica Systems) using a 40X (1.2 NA) or 63X (1.4 NA) oil-immersion lens. Whole-mount immunostained embryos were imaged using a 25X (0.95 NA) water-immersion lens. All confocal images were processed using Volocity (Improvision) or Imaris software and all figures were composed with Adobe Photoshop and Adobe Illustrator.

### Quantitative RT-PCR

Mutant and sibling zebrafish embryos were sorted by phenotype at 48hpf. Total RNA was isolated using Trizol according to manufacturer’s instructions. cDNA was synthesized and amplified with Transplex Whole Transcriptome Amplification Kit (Sigma) using 50 nanograms of total RNA, according to the protocol provided. Nucleic acid concentrations were obtained using a Nanodrop. Real-time PCR experiments were performed in triplicate using GoTaq qPCR Master Mix (Promega) in a BioRad iCycler. Fold changes in RNA levels were calculated according to the ΔΔCt method [80], and expression was normalized to β-actin levels. Quantitect primers (Qiagen) were used to amplify *aldh1a3* (QT02111613), *actb1* (QT02174907), *ccnd1* (QT02178519), *chd7* (QT02068584), *col10a1a* (QT02129708), *crabp1b* (QT02229297), *elavl2* (QT02092713), *gnb3a* (QT02129701), *h2afx* (QT02140278), *hat1* (QT02209501), *hdac1* (QT02157099), *ntn1a* (QT02422490), *nr2f5* (QT02125424), *rx1* (QT02067786), *shha* (QT02236136), *shhab* (QT02055312), *smarcc2* (QT02101008), *tdrd6* (QT02235821), *tfap2* (QT02211622), and *vsx2* (QT02050426).

## Acknowledgements

We are very grateful to members of our group for suggestions and critically reading the manuscript. We particularly thank Eirinn Mackay for his technical assistance. We thank Carole Wilson and her team for fish care. This work was funded by grants from the Medical Research Council [G0900994 and MR/L003775/1 to S.W.W. and G.G] and Wellcome Trust [095722/Z/11/Z and 207483/Z/17/Z to R.J.P, and 089227/Z09/Z and 104682/Z/14/Z to S.W.W. and FONDECYT [11160951 to L.E.V].

## Author contributions

GG, KT, JH and SWW conceived and planned experiments and wrote the paper with help from RJP; GG, KT, JH, LV, KC, WF conducted experiments; GR and RG contributed to initial mapping of the mutation which was refined by JH and RJP who devised the SNP mapping approach; CH participated in the genetic screen to identify the eye and midline defects; MR helped to supervise research undertaken by JH.

---

**Figure S1.**
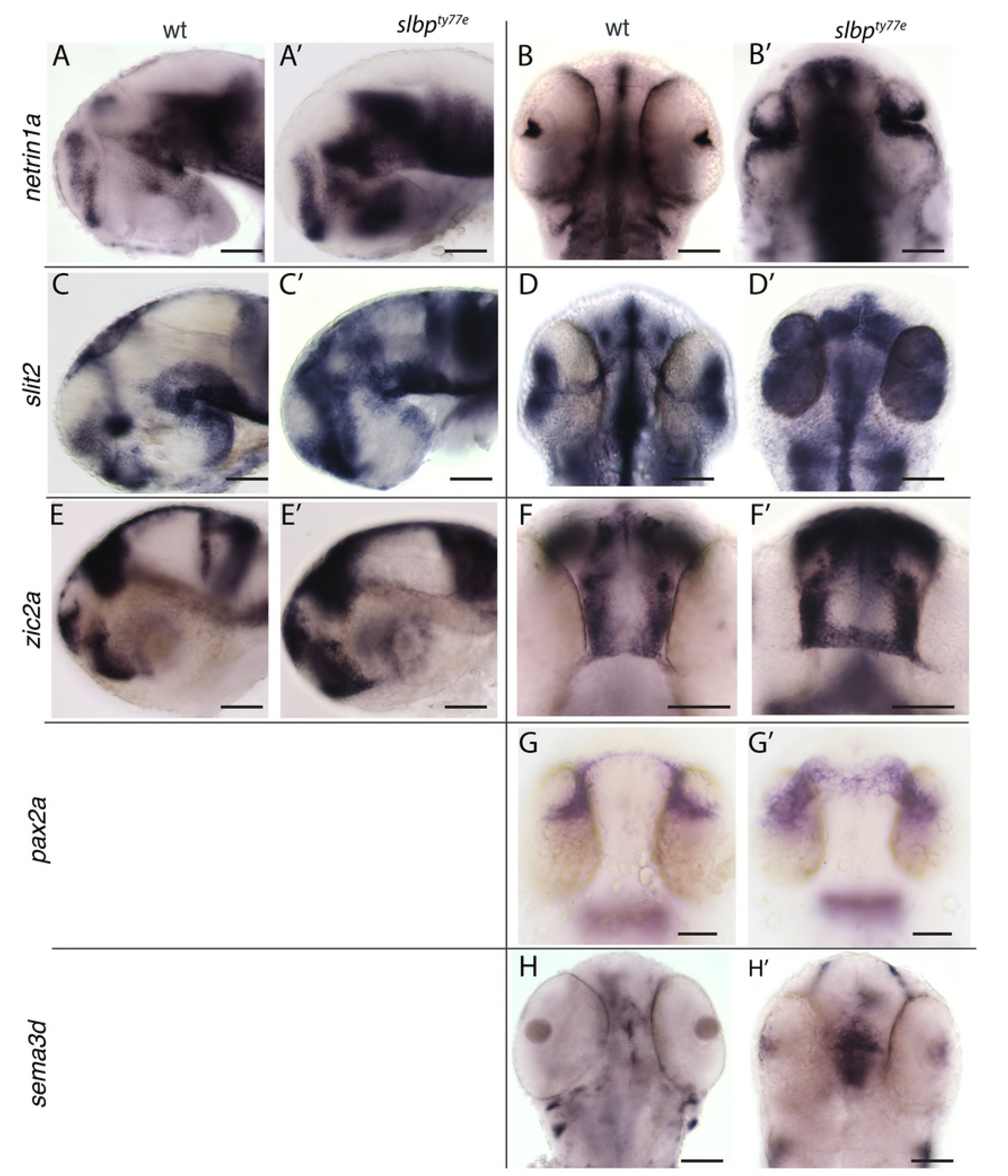
Misregulation of expression of genes potentially implicated in midline axon guidance in *slbp^ty77e^* mutants. Views of heads/brains of wildtype (A-H) and *slbp*^ty77e^ (A’-H’) embryos showing expression of genes indicated to the left of each row. Genotype is indicated at top of each column. Lateral views (A,A’,C,C’,E,E’); dorsal views (B,B’,D,D’,F,F’,G,G’,H,H’). All embryos are 60hpf apart from G,G’ which are 30hpf. Scale bars: 100μm.

**Figure S2.**
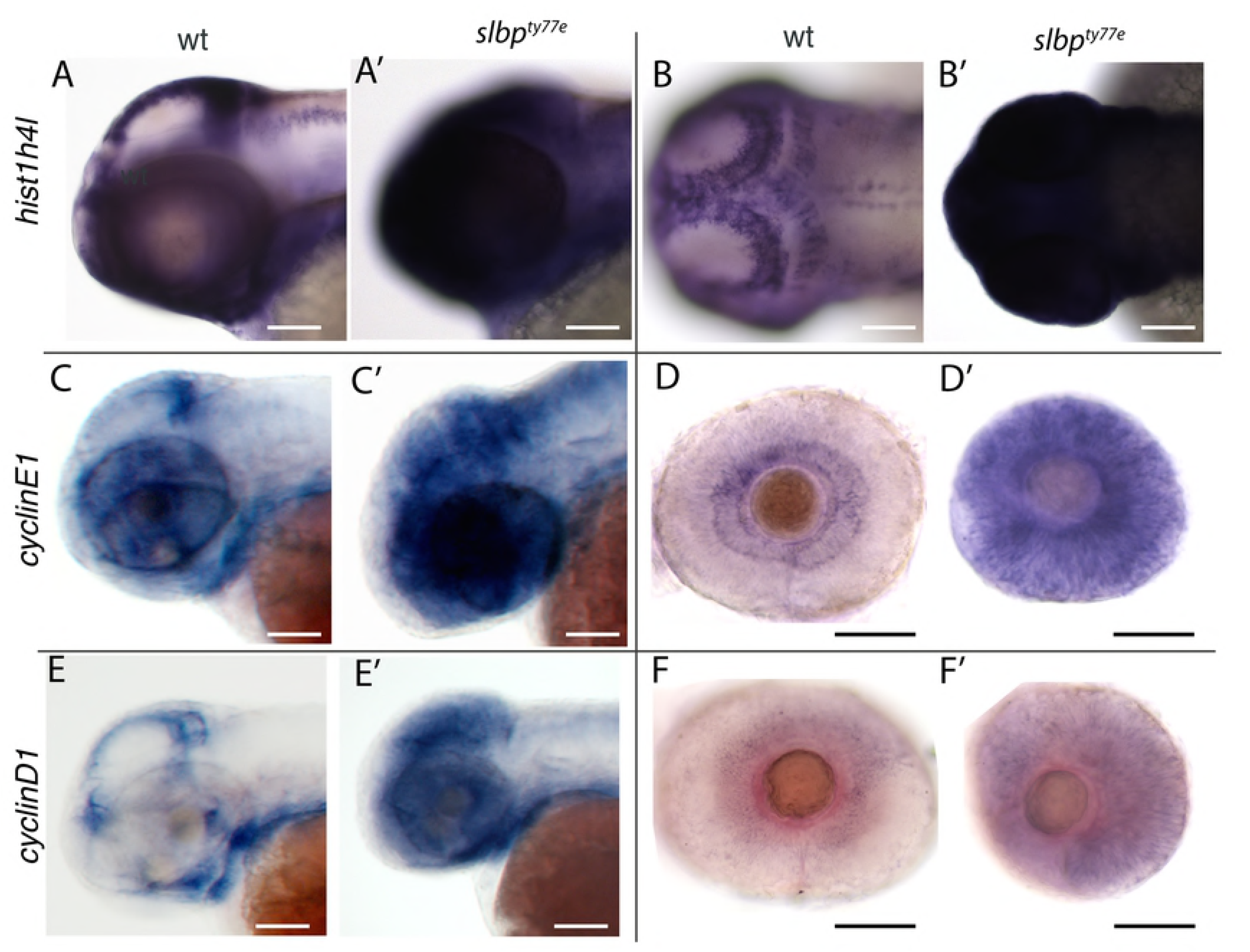
Upregulation of cell cycle markers in *slbp^ty77e^* mutants. Images of wildtype (wt, A-F) and *slbp^ty77e^* (A’-F’) heads (A-C’; E,E’) and eyes (D-F’) at 60hpf showing expression of genes indicated to the left of each row. Genotypes indicated at top of each column. Lateral (A,A’; C-F’) and dorsal view (B,B’). Scale bars: 100μm.

**Table S1. List of transcripts with differential expression between wildtype and *slbp^ty77e^* mutants.**

Unprocessed transcript list derived from the differential expression analysis performed on the BAM files from all three biological replicates and the merged transcript dataset using Cuffdiff.

**Table S2. Gene list used for GO term enrichment analysis for “Biological Process” on all of the upregulated genes showing a significant change in expression (q value ≤ 0.01) in our RNAseq data.**

(Sheet 1) Upregulated genes sorted by q value.

(Sheet 2). Upregulated genes sorted by log2(fold change).

(Sheet 3) List of GO terms related to “Biological Process” generated using the AmiGO2 tool (The Gene Ontology Consortium) manually grouped into 14 categories (Listed in Fig 6B).

(Sheet 4) Manual categories used to generate the GO term pie chart in Figure 6B’.

**Table S3. Gene list used for GO term enrichment analysis for “Biological Process” on all of the downregulated genes showing a significant change in expression (q value ≤ 0.01) in our RNAseq data.**

(Sheet 1) Downregulated genes sorted by q value.

(Sheet 2). Downregulated genes sorted by log2(fold change).

(Sheet 3) List of GO terms related to “Biological Process” generated using the AmiGO2 tool (The Gene Ontology Consortium) manually grouped into 14 categories (Listed in Fig 6B).

(Sheet 4) Manual categories used to generate the GO term pie chart in Figure 6B’’.

**Table S4. Manually curated list of genes showing significant changes in expression level related to nervous system development, cell cycle and histones.**

(Sheet 1) Downregulated genes with a log2(fold change >-2) related to neural Development, axon pathfinding and synaptogenesis.

(Sheet 2) Upregulated genes related to cell cycle.

(Sheet 3) Histone related genes all show a log2(fold change >2.5). Histone subunit genes enriched in our dataset are largely found in two chromosomal regions on chromosome 7 and chromosome 25.

